# Mutations in the spike RBD of SARS-CoV-2 omicron variant may increase infectivity without dramatically altering the efficacy of current multi-dosage vaccinations

**DOI:** 10.1101/2021.12.08.471688

**Authors:** Bingrui Li, Xin Lu, Kathleen M. McAndrews, Raghu Kalluri

## Abstract

With the continuous evolution of SARS-CoV-2, variants of concern (VOCs) and their mutations are a focus of rapid assessment. Vital mutations in the VOC are found in spike protein, particularly in the receptor binding domain (RBD), which directly interacts with ACE2 on the host cell membrane, a key determinant of the binding affinity and cell entry. With the reporting of the most recent VOC, omicron, we performed amino acid sequence alignment of the omicron spike protein with that of the wild type and other VOCs. Although it shares several conserved mutations with other variants, we found that omicron has a large number of unique mutations. We applied the Hopp-Woods scale to calculate the hydrophilicity scores of the amino acid stretches of the RBD and the entire spike protein, and found 3 new hydrophilic regions in the RBD of omicron, implying exposure to water, with the potential to bind proteins such as ACE2 increasing transmissibility and infectivity. However, careful analysis reveals that most of the exposed domains of spike protein can serve as antigenic epitopes for generating B cell and T cell-mediated immune responses. This suggests that in the collection of polyclonal antibodies to various epitopes generated after multiple doses of vaccination, some can likely still bind to the omicron spike protein and the RBD to prevent severe clinical disease. In summary, while the omicron variant might result in more infectivity, it can still bind to a reasonable repertoire of antibodies generated by multiple doses of current vaccines likely preventing severe disease. Effective vaccines may not universally prevent opportunistic infections but can prevent the sequelae of severe disease, as observed for the delta variant. This might still be the case with the omicron variant, albeit, with increased frequency of infection.

## Introduction

The causative virus of the COVID-19 pandemic, SARS-CoV-2, continues to evolve into new variants since it was first identified. Even though vaccines have been widely administered, some of the emerging variants induced putatively lower immunogenicity causing decreased protection from infection by current vaccines (Collier, De Marco et al. 2021) (Bian, Gao et al. 2021). These variants were listed as variants of concern (VOCs) with increased transmissibility and infectivity, and reduced antigenicity (Harvey, Carabelli et al. 2021), requiring further investigation for the understanding of viral properties and therapeutics development. The most recently identified variant, omicron, reported on November 24^th^, 2021, was found to have many additional mutations compared with previous variants, further highlighting the urgency of such studies.

Spike protein (S-protein) mediates the interaction between the virus and the receptor angiotensin-converting enzyme 2 (ACE2) on the host cell membrane and is the major target for immunogenicity (Tortorici and Veesler 2019). The receptor-binding domain (RBD) is the region directly interacting with ACE2 which makes it the crucial determinant of host specificity (Wang, Zhang et al. 2020). Most of the key mutations postulated to increase the spread of the virus also occurred in RBD.

While further analysis and unraveling of the S-protein structure and properties of omicron is underway, understanding of how the omicron variant may change the binding affinity with ACE2 and influence the current vaccination effect is under urgent need. Computational analysis and predictions based on the viral genome and amino acid sequences provide a fast and powerful tool to investigate the mutations occurring in the variants and reveal the potential impact of these mutations on viral properties.

One main element affecting the infectivity and transmissibility of the virus is the binding affinity between the ligand (S-protein) and the receptor (ACE2) (Liu, Zhang et al. 2021), which is largely determined by the interaction at the molecular level and the formation of molecular bonds (Chanphai, Bekale et al. 2015). With the data from current mutations and variants, the binding affinity between S-protein and ACE2 in the variants compared with that in the wide type has been shown to be enhanced partially by the increased hydrophilicity which is due to the change of the electrostatic and the van der Waals energies caused by the formation of extra hydrogen-bonds and salt bridges (Han, Wang et al. 2021) (Ali, Kasry et al. 2021). The Hopp-Woods hydrophobicity scale is an algorithm to calculate the hydrophobicity index of the amino acid residues where polar residues have been assigned positive values and nonpolar residues negative values. It is primarily designed to find potentially antigenic sites in proteins (Hopp 1989).

In this study, we compared the protein sequence of the major VOCs with the reference SARS-CoV-2 virus and studied alterations in the hydrophobicity of the amino acid residues of RBD in omicron. The predicted hydrophilicity of RBD for different variants was generated, giving insight into how transmissibility and infectivity of the variant may change and providing knowledge for the potential understanding of the omicron variant.

## Methods

### Genomic and protein sequence analysis

Analyses were performed to identify and compare the genomic mutations of SARS-CoV-2 variants. We downloaded the complete genomes of the original reference (EPI_ISL_402124) and five emerging variants, including alpha, beta, gamma, delta and omicron, from the GISAID (https://www.gisaid.org/) database. For the variant sequences, we selected one complete genome (>29□000□bp) with high coverage that was earliest detected in the location where the variant was first reported. The genome sequences were translated to protein sequences using the Expasy Translate tool (https://web.expasy.org/translate/). The regions of spike protein and RBD were identified based on the genome locations. The multiple sequence alignment of the protein sequence was carried out by Clustal Omega (https://www.ebi.ac.uk/Tools/msa/clustalo/) with the default parameters. The multiple sequence alignment plots were generated by ggmsa (v1.1.2) (Zhou and Yu 2021).

### Hydrophilicity/Hydrophobicity analysis

The hydrophilicity/hydrophobicity analysis was performed by ProtScale (https://web.expasy.org/protscale/) of both spike protein and RBD regions using Hopp & Woods amino acid scale with the window size of 5. The plots were generated by the ggplot2 (v3.3.5) package implemented in R (v4.1.2).

## Results

### Comprehensive amino acid mutation analyses reveal the distinct patterns in the RBD region of five SARS-CoV-2 variants

We obtained the complete genomes of five SARS-CoV-2 VOC, alpha (B.1.1.7), beta (B.1.351), gamma (P.1), delta (B.1.617.2) and omicron (B.1.1.529), and the original reference genome (EPI_ISL_402124) from the GISAID (https://www.gisaid.org/) database. For each variant, we selected one complete genome sequence with high coverage detected in the location where the variant was first reported in the database. For example, we downloaded the genome sequence of the omicron (EPI_ISL_6913995) variant that was collected on November 18th, 2021 submitted by the lab in South Africa. The amino acid sequence of spike protein and RBD region were extracted based on the sites widely reported in previous studies. Afterwards, we translated the genome sequence to amino acid sequence using the Expasy Translate tool (https://web.expasy.org/translate/).

We aligned the amino acid sequences of RBD of the five SARS-CoV-2 variants to that of the reference separately (**Figure 1**). In general, no amino acid insertions or deletions were found within the RBD of the five variants, while distinct amino acid mutations were detected from each variant, especially the variant omicron. Specifically, N501Y is the only variation found in RBD of the variant alpha (**Figure 1A**), while this mutation commonly exists in all the other variants except for the variant delta. Three mutations of amino acids were detected in the RBD of the variant beta, including K417N, E484K and N501Y (**Figure 1B**). Of note, the mutation K417N was found exclusively in the variants beta and omicron, whereas the mutation in the same site was found in the variant gamma, but lysine (K) was substituted by threonine (T) instead of asparagine (N) (**Figure 1C**). As for the variant delta, mutations L452R and T478K were found in RBD (**Figure 1D**), and L457R was the unique mutation among all the five variants. The mutation T478K was found in both the variants delta and omicron. Obviously, the variant delta displayed quite unique mutation patterns compared to the previously reported variants. Given that the variant delta was believed to be more than twice as contagious as the previous variants (Twohig, Katherine A et al. 2021), these two distinct mutations may play important roles in the increased infectivity and transmissibility. Recently, the emergence of the variant omicron raised a large number of concerns for the community due to a total of 15 amino acid mutations found in RBD, along with one common mutation with delta. In addition to the four common mutations identified in other variants (K417N, T478K, E484A and N501Y), the omicron variant possesses 11 distinct mutations in RBD, including G339D, S371L, S373P, S375F, N440X, G446S, S477N, Q493R, G496S, Q498R and Y505H (**Figure 1E, Figure 2**).

**Figure 1.**
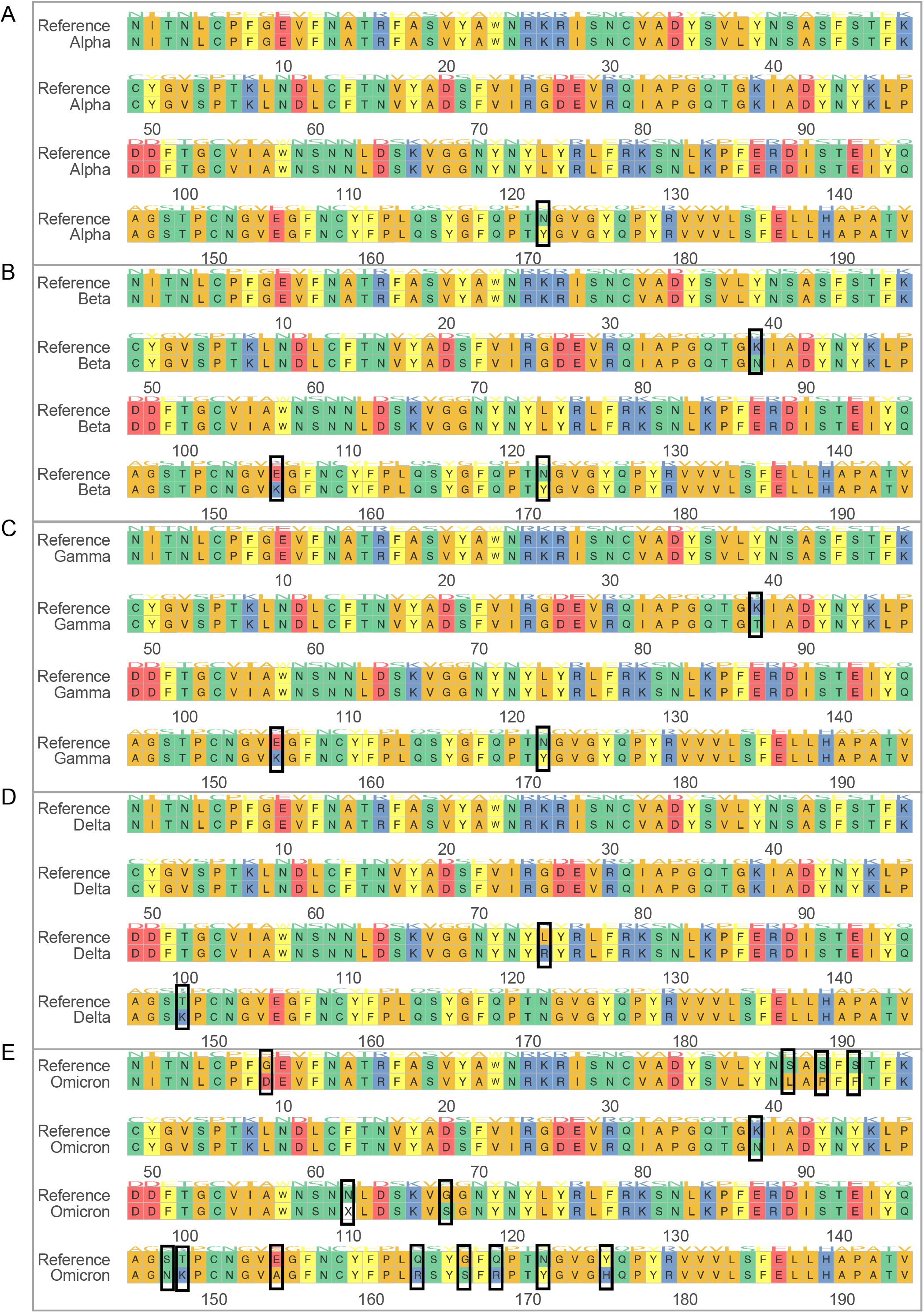
Multiple sequence alignment of amino acids between each variant and reference. (**A**) Alpha variant vs. reference. (**B**) Beta variant vs. reference. (**C**) Gamma vs. reference. (**D**) Delta vs. reference. (**E**) Omicron vs. reference. Black boxes denote residues with mutations.

**Figure 2.**
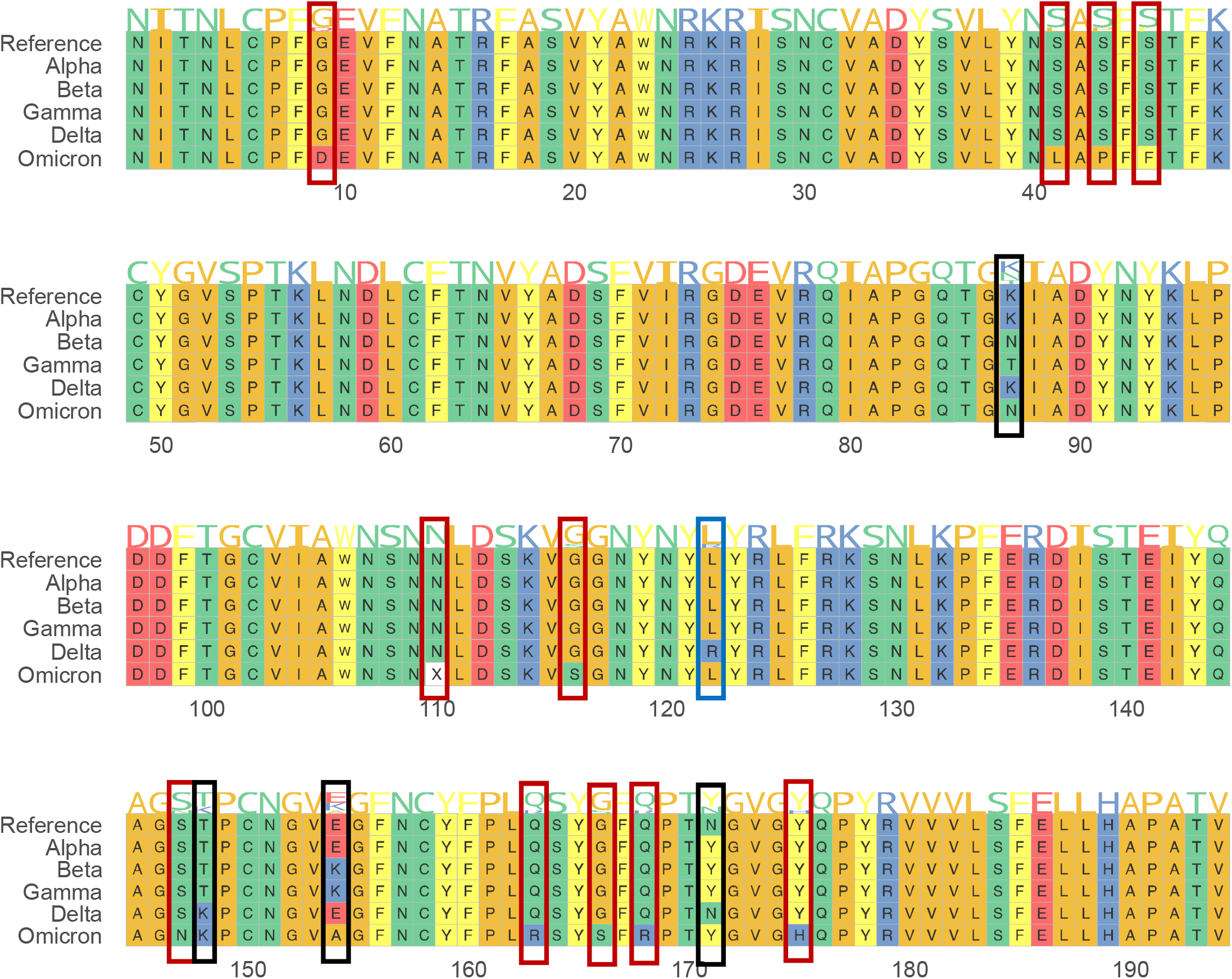
Multiple sequence alignment of amino acids between variants and reference. Red boxes denote mutations unique to omicron, blue boxes denote mutations unique to delta, and black boxes denote mutations detected in multiple variants.

Although the site of mutation was shared by three of the previous variants, omicron displayed a distinct substitution from glutamic acid (E) to alanine (A) instead of from glutamic acid (E) to lysine (K) in both the variants beta and gamma (amino acid 157 of RBD, **Figure 2**). Furthermore, compared to the number of mutations identified in the previous variants, there is a significantly increased number of mutations in omicron, which may suggest potentially increased transmissibility and reduced effects of the neutralizing antibodies.

To further investigate the impact of mutations on the binding affinity between SARS-CoV-2 and ACE2, we obtained the contacting residues from previous studies (Lan, Jun, et al 2020). We examined the sites of these residues in our multiple alignment results and labeled the mutations occurring in different variants (**Figure 3**). As expected, Omicron has five unique mutations out of the 17 contacting residues in the RBD region.

**Figure 3.**
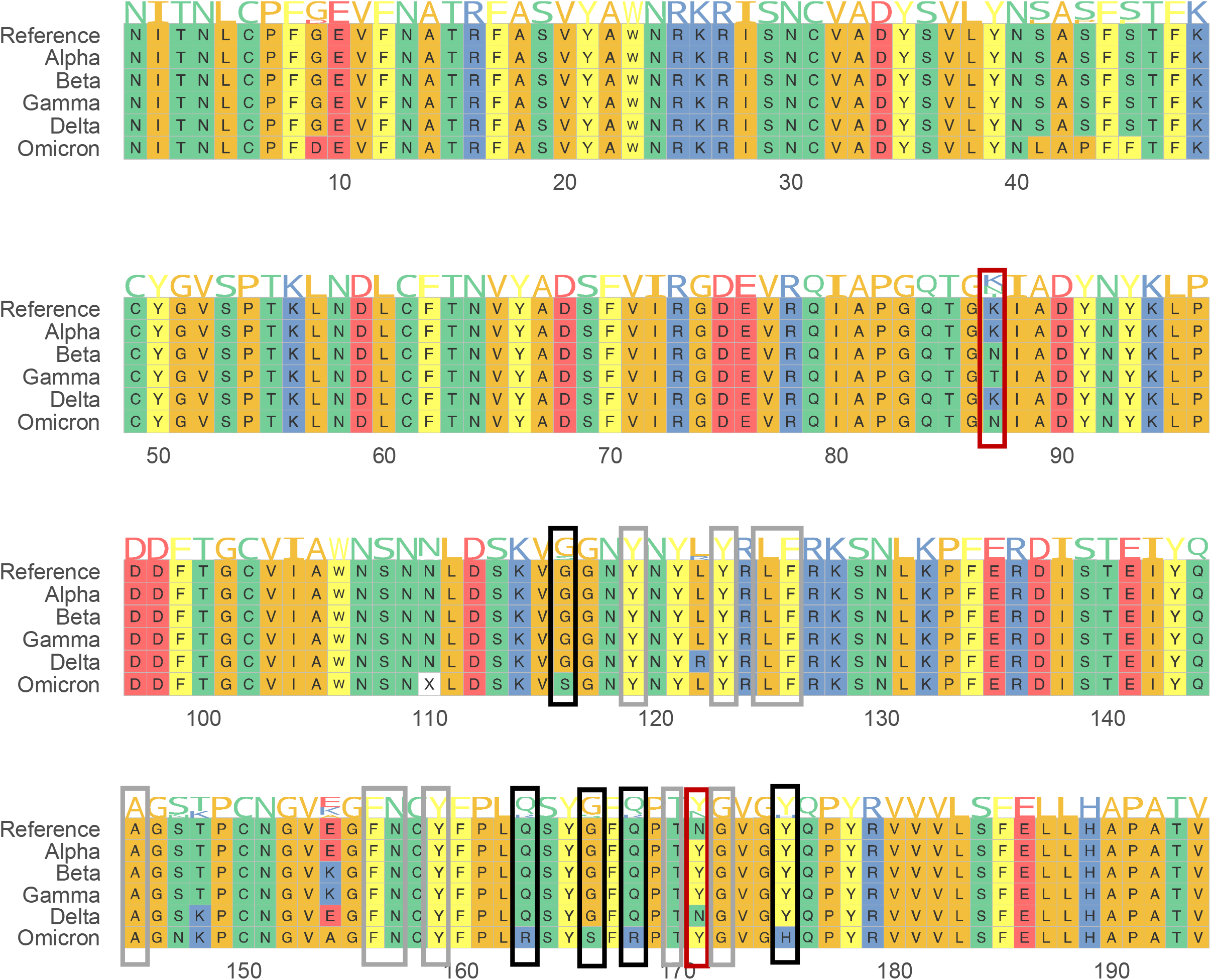
Contacting residues between SARS-CoV-2 and ACE2. Boxes denote the contacting residues. Black boxes denote mutations unique to omicron, red boxes denote mutations occurring in multiple variants, and grey boxes denote no mutations in any variant.

### Hydrophilicity analysis shows increased hydrophilicity caused by mutations

To further investigate the potential impact of mutations in the virus properties, we performed hydrophilicity analysis on the RBD of each variant based on the Hood-Woods scale and compared them with the reference genome (**Figure 4**). Interestingly, compared to the mutations occurred in alpha, beta and gamma, the two mutations detected in delta increased the hydrophilicity obviously and even reversed the mutation region from hydrophobicity to hydrophilicity (**Figure 4B-E**), which may strengthen the binding affinity between RBD and ACE2 and thus increase the transmissibility of the virus. As for omicron, in addition to the T478K mutation which changed the region from hydrophobicity to hydrophilicity, G339D caused a similar pattern changing the region from hydrophobicity to hydrophilicity. Q493R, G496S and Q498R mutations detected in omicron were also shown to increase the hydrophilicity of the particular regions surrounding the mutations (**Figure 3E**). Overall, the mutations detected in the variants showed significant impacts on the hydrophilicity/hydrophobicity properties, which may help explain the distinct properties of variants in terms of their transmissibility and pathogenicity (**Figure 4**).

**Figure 4.**
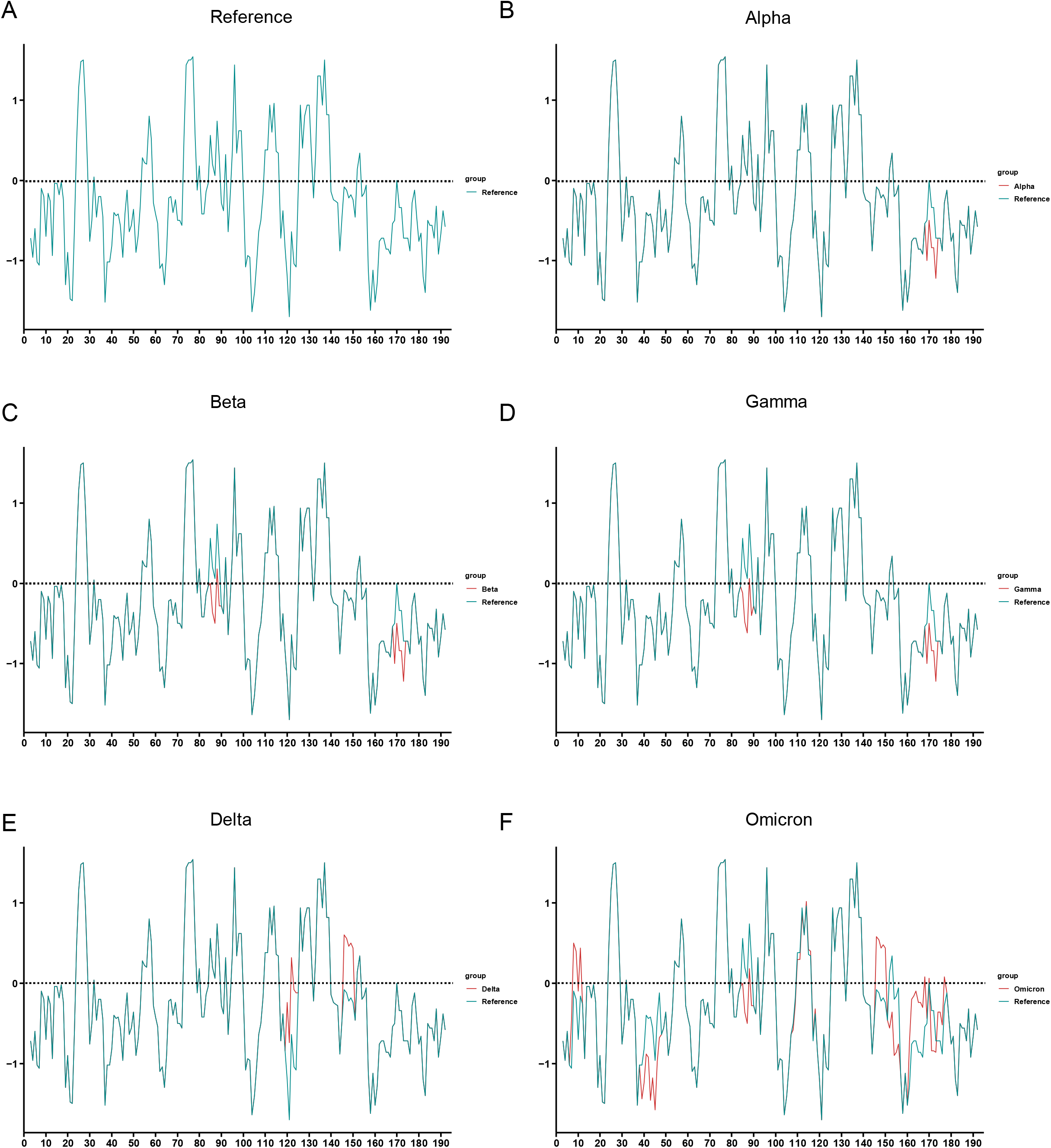
Hydrophilicity plots of each variant and reference in the RBD domain. Comparisons of hydrophilicity for (**A**) Reference. (B) Alpha variant vs. reference. (**C**) Beta variant vs. reference. (**D**) Gamma vs. reference. (**E**) Delta vs. reference. (**F**) Omicron vs. reference.

**Figure 5.**
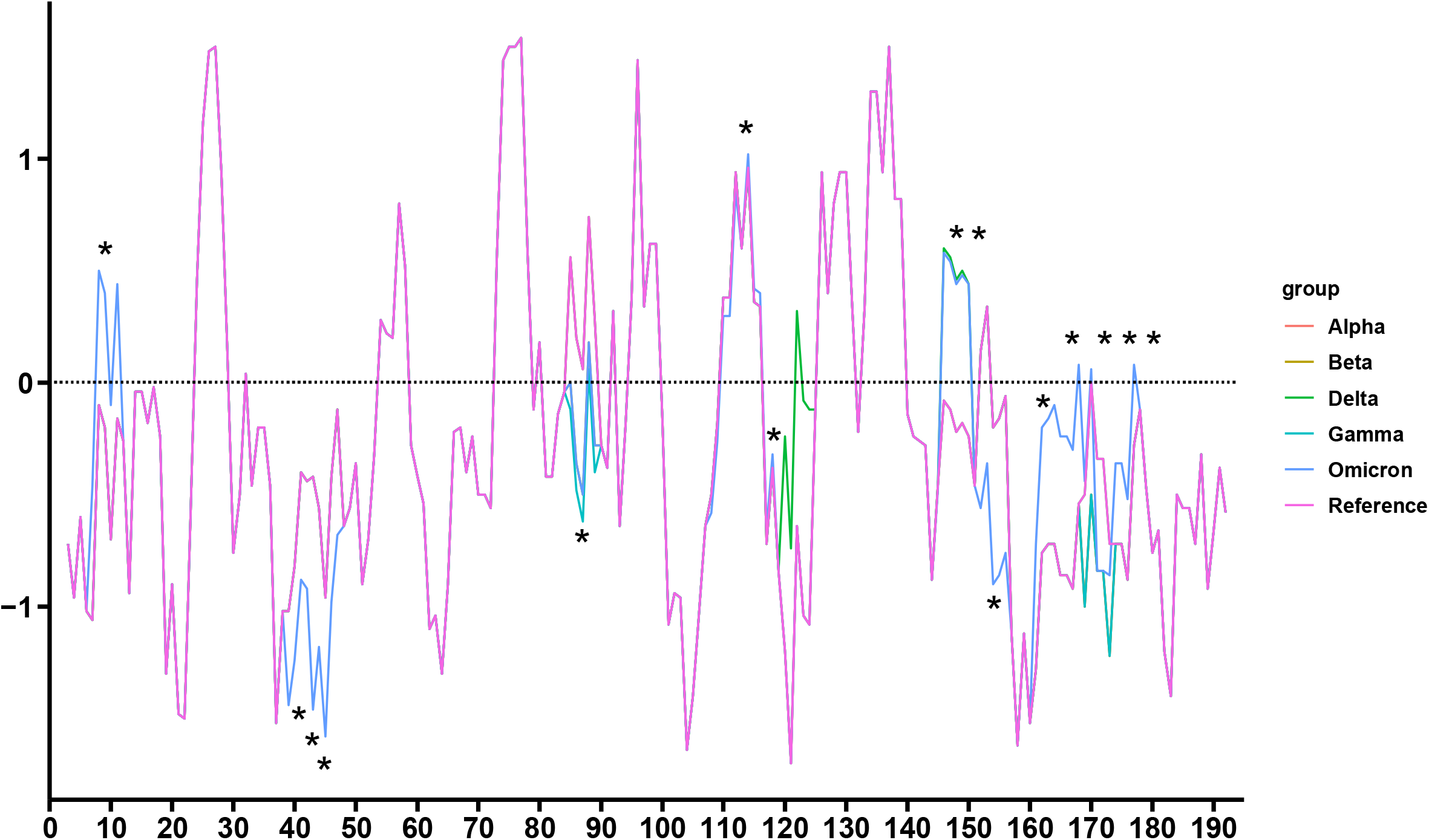
Hydrophilicity plots of all variants and the reference. Asterisks denote mutations unique to omicron.

### Amino acid sequence analyses of spike protein display a significant increase in the number of mutations of the omicron variant

We next sought to investigate the spike protein sequences of delta and omicron due to the similarity in the mutation patterns and potential impacts on public health. In addition to the substitutions detected in RBD, we found four additional and thirteen unique substitutions in delta and omicron, respectively. One deletion (sites 156-157) was found in N-Terminal Domain (NTD) of delta, while two deletions (sites 69-70; sites 143-145) and one insertion (sites 213-214) were identified in NTD of omicron (**Figure 6**). There were no common mutations detected in NTD of delta and omicron, while two substitutions (sites 614 and 681) were found at the same sites in subdomain 2 (SD2). At site 614, the aspartic acid was substituted for glycine for both delta and omicron, whereas proline (P) was substituted by arginine (R) and histidine (H) in delta and omicron, respectively. In total, delta and omicron have 9 and 34 mutations, respectively.

**Figure 6.**
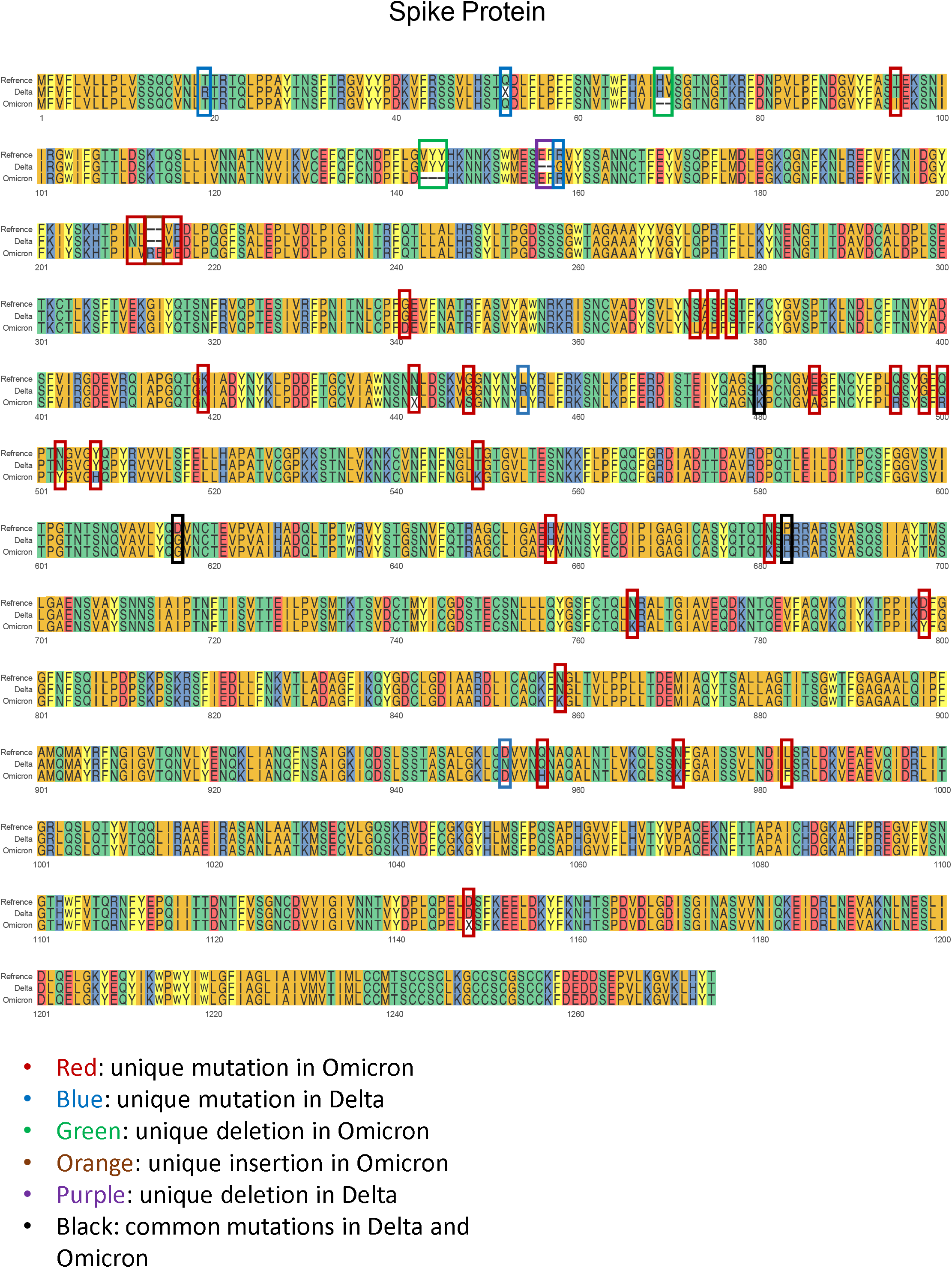
Multiple sequence alignment of amino acids for the spike proteins of reference, Delta and Omicron. Red boxes denote mutations unique to omicron, blue boxes denote mutations unique to delta, green boxes denote deletions unique to omicron, orange boxes denote insertions unique to omicron, purple boxes denote deletions unique to delta, and black boxes denote mutations common to delta and omicron.

### Hydrophilicity analyses of the entire spike protein imply the impact of mutations on the hydrophilicity and hydrophobicity of the proteins

To reveal the impact of mutations on the hydrophilicity and hydrophobicity for the entire spike protein, we performed hydrophilicity analyses of the spike amino acid sequences of the variants delta, omicron, and the reference and compared the hydrophilicity score between each variant (**Figure 7**). We first compared the hydrophilicity score between delta and the reference (**Figure 8**). We observed several hydrophilicity peaks caused by the mutations in the variant delta. For example, T19R dramatically increased the hydrophilicity score surrounding site 19. Likewise, we compared the hydrophilicity score between omicron and the reference and found multiple hydrophilicity peaks caused by the mutations (**Figure 8B**). Obviously, there were more mutation-caused hydrophilicity peaks in omicron compared to delta. Furthermore, we compared omicron and delta in terms of the hydrophilicity score and the results confirmed that the mutations identified in omicron increased the hydrophilicity score compared to that of delta (**Figure 8C**).

**Figure 7.**
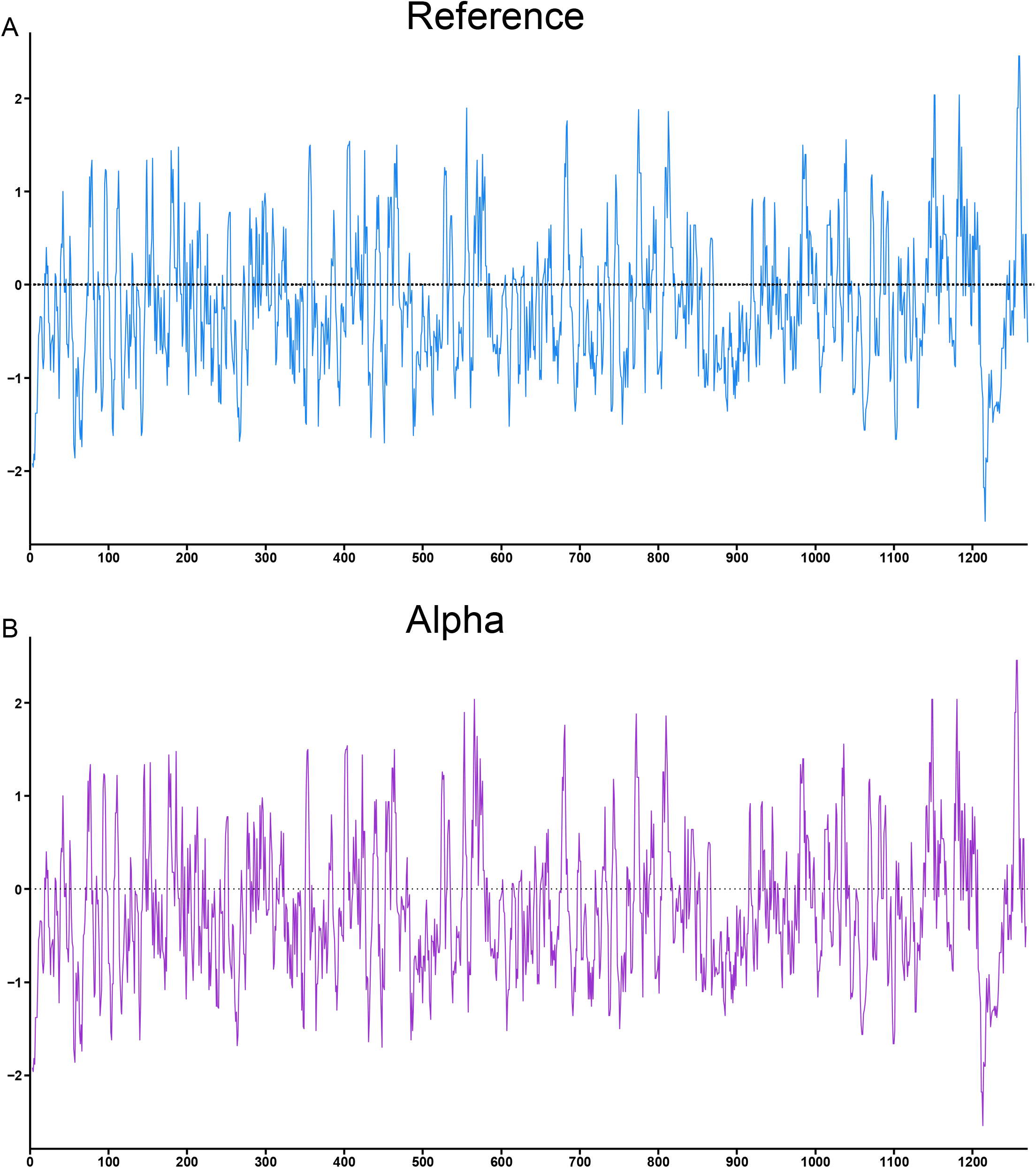
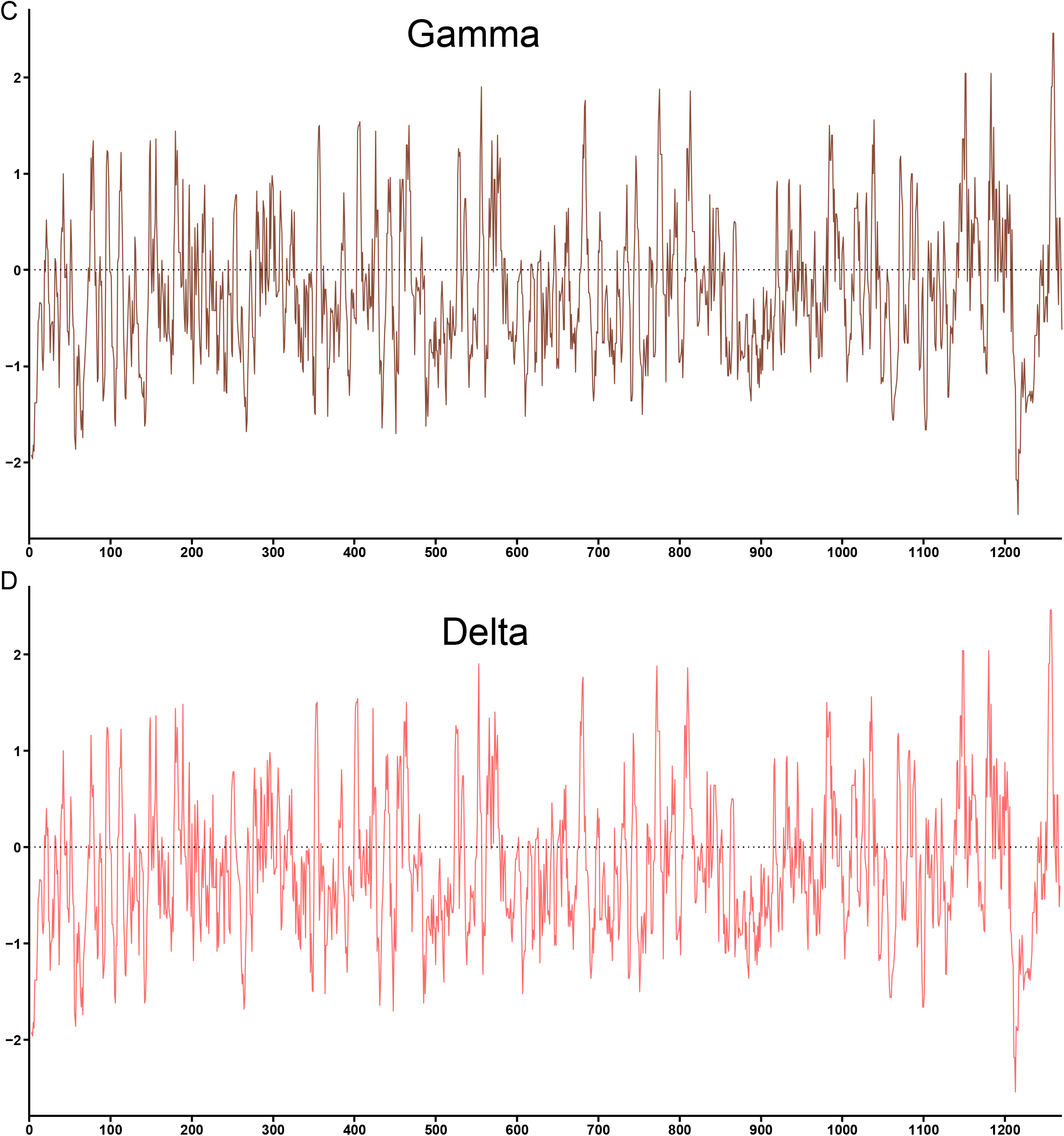
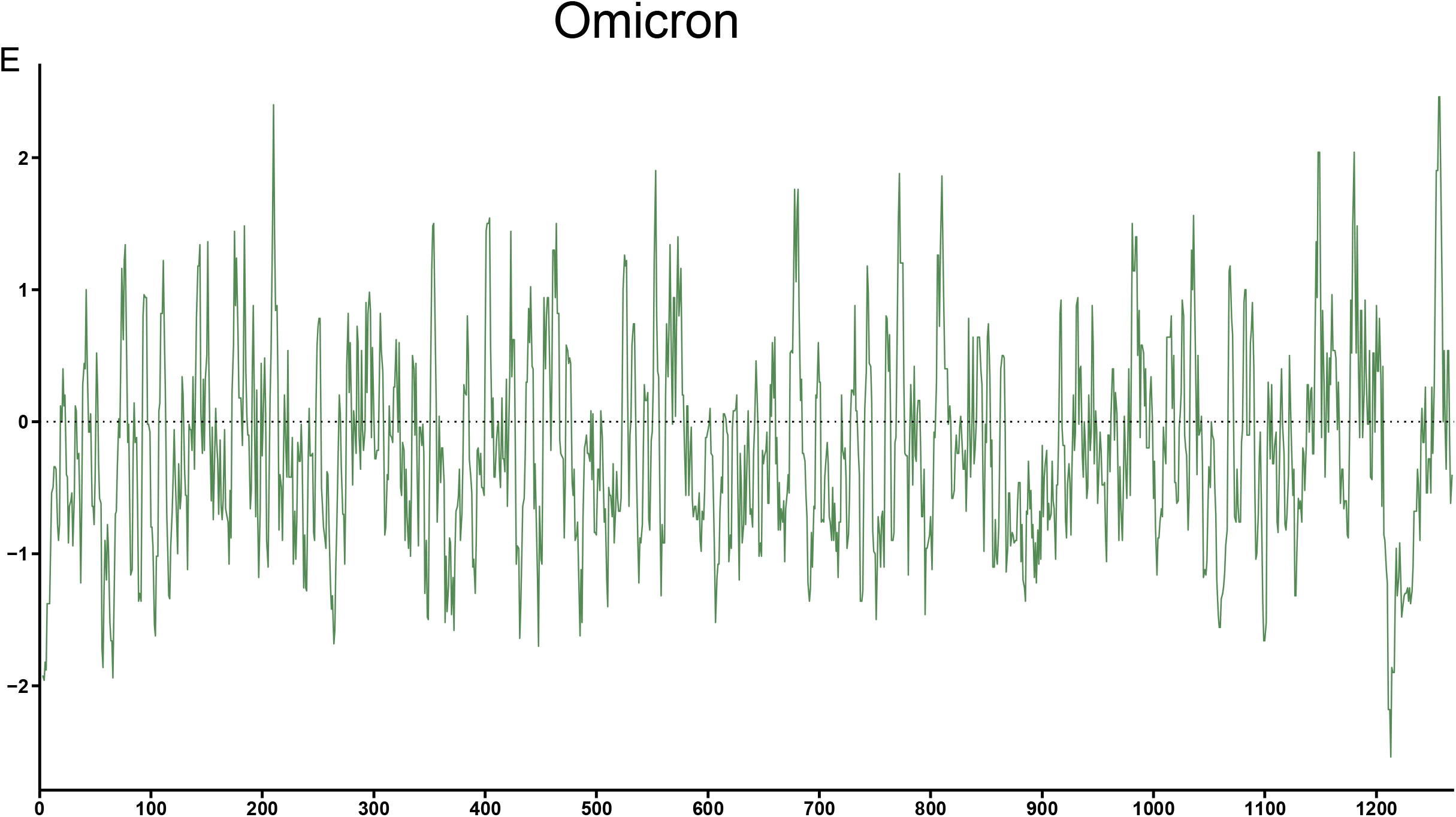
Hydrophilicity plots of spike proteins for all variants. **(A**) Reference. (**B**) Alpha. (**C**) Gamma. (**D**) Delta. (**E**) Omicron.

**Figure 8.**
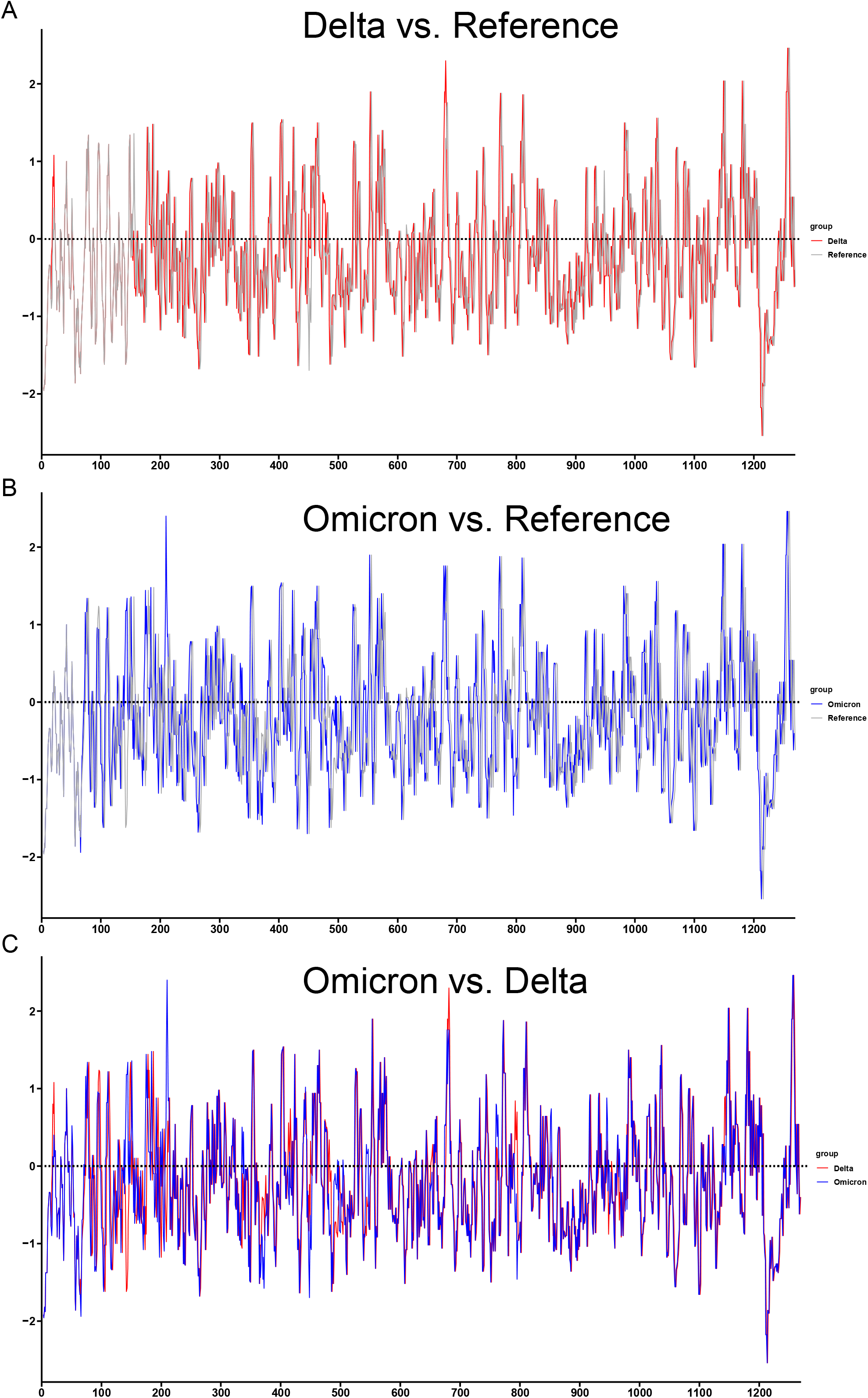
Comparisons of hydrophilicity scores of spike proteins for reference, Delta and Omicron. Comparisons of hydrophilicity scores for (**A**) Delta vs. reference. (**B**) Omicron vs. reference. (**C**) Omicron vs. Delta.

## Conclusion and Discussion

In this study, we compared the mutations of the major VOCs of SARS-CoV-2 and studied the hydrophobicity of the amino acid residues induced by the variants based on the Hopp-Woods hydrophobicity scale. Our findings associate a large number of novel mutations in the RBD of omicron and several regions with increased hydrophilicity, potentially suggestive of enhanced binding affinity between the RBD of omicron variant and ACE2, which may increase the contagiousness of omicron. Our study provides evidence of potential changes in transmissibility and infectivity of the new variant and the need for further evaluation of the spike protein structure.

The relationship between hydrophobicity of the amino acid residues and the binding affinity between S-protein and ACE2, however, is not absolute and determined simply from the analysis of the amino acid residues. It is also influenced by higher-order protein structure and other parameters. There are several regions in RBD of omicron displaying higher hydrophobicity compared with that of the reference virus, which could influence the binding affinity positively in some positive ways. More information from the protein is needed to better predict and understand the transmission and infection patterns of the variants.

## Acknowledgements

This research in the R.K. lab was supported by Lyda Hill Philanthropies^®^.

## Author contributions

Conceptualization: R.K.; formal analysis: B.L., X.L.; writing – original draft: B.L., X.L.; writing – review & editing: B.L., X.L., K.M., R.K.

## Notes

### Competing Interest Statement

The authors have declared no competing interest.

